# A folder mechanism ensures size uniformity among *C. elegans* individuals by coupling growth and development

**DOI:** 10.1101/2021.03.24.436858

**Authors:** Benjamin D. Towbin, Helge Grosshans

## Abstract

Animals increase by orders of magnitude in their volume during development. Hence, even small differences in the growth rates between individuals could generate large differences in their adult body size. Yet, such volume divergence among individuals is usually not observed in nature.

We combined theory and experiment to understand the mechanisms of body size uniformity. Using live imaging, we measured the volume growth of hundreds of individuals of *C. elegans* over the entire span of their postembryonic development. We find that *C. elegans* grows exponentially in volume with a coefficient of variation of the growth rate of ∼7%, but that individuals diverge much less in volume than expected from this heterogeneity. The mechanism counteracting size divergence does not involve size thresholds for developmental milestones. Instead, an inverse coupling of the growth rate and the duration of development produces a constant volume fold change per larval stage.

The duration of larval stages of *C. elegans* is determined by the period of a developmental oscillator. Using mathematical modelling, we show that an anti-correlation between the growth rate and the oscillatory period emerges as an intrinsic property of a genetic oscillator. We propose that the robustness of body volume fold change is a hard-wired characteristic of the oscillatory circuit and does not require elaborate mechanisms of size control by cellular signalling. Indeed, the coupling of growth and development was unaltered by mutation of canonical pathways of growth control. This novel concept of size homeostasis may broadly apply to other multicellular systems controlled by genetic oscillators.

## Introduction

Animals increase by orders of magnitude in volume during development. Despite this massive increase, individuals of the same species differ only little in their adult body volume, although even small differences in the growth rate could, in principle, amplify to large differences in size (Fig. 1a). Indeed, growth to the appropriate size is crucial for animal fitness and under strong selective pressure ^1^.

**Figure 1.**
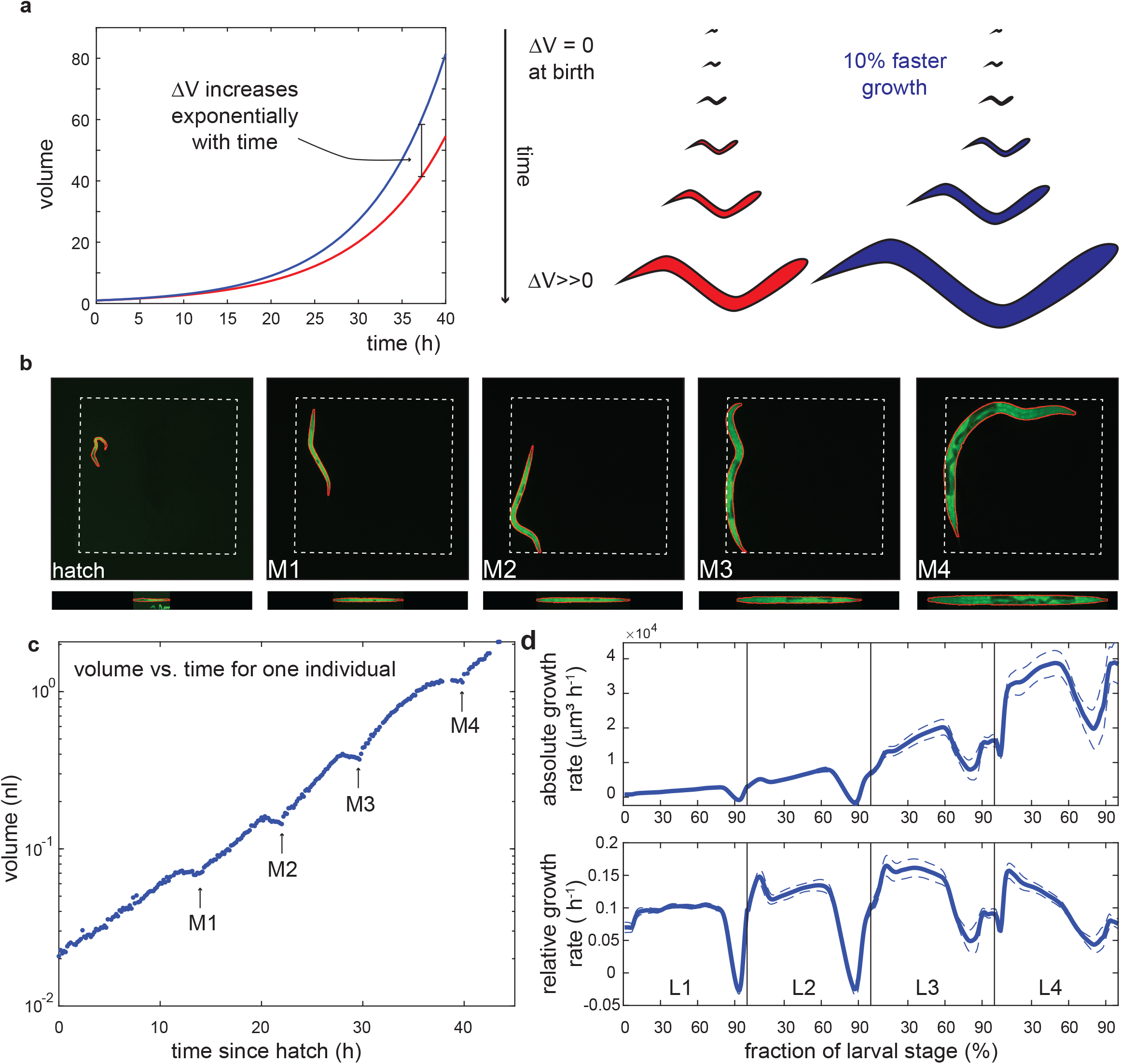
*C. elegans* grows faster than linearly within larval stages. *(related Supplemental Figure S1)* a. Illustration of volume divergence during exponential growth. Blue individual has a 10% faster growth rate than red individual (0.1/h vs. 0.11/h). b. Images of the same individual at birth and moult 1 (M1) to moult 4 (M4). Dotted square indicates edge of the chamber with dimensions of 600 µm x 600 µm. The red line shows the segmentation outline. Images at the bottom are computationally straightened animals used for volume computation. C. Volume measurement of one individual as a function of time starting at hatching. Arrows indicate larval stage transitions detected as re-start of growth after lethargus. d. Average absolute (µ_abs_) and relative (µ_rel_) growth rate during development. The x-axis indicates the fraction τ of larval stage progression (τ = [time since last moult]/[total duration of larval stage]). Average growth rates were computed by re-scaling individuals according to moulting times and averaging growth rates across all individuals at the same τ. Dips in the growth rate correspond to growth halt during lethargus. Dashed lines are 95% confidence intervals for the mean of day-to-day repeats.

Although molecular pathways that promote or limit the growth of cells, organs, or organisms have been characterized extensively ^2,3^, the mechanisms responsible for size uniformity among individuals are poorly understood. Studies of animal size control in the context of environmental and genetic perturbations ^2,3^ led to the hypothesis that specific size thresholds are associated with the passing of developmental milestones. For example, insects commit to enter metamorphosis at a critical weight ^4^. Similarly, the body volume of *C. elegans* at larval stage transitions is nearly invariant under dietary restriction ^5^, and growth retardation of humans due to malnutrition or hormonal imbalance is compensated by catch-up growth later in life ^6,7^. However, measuring the response to exogenous perturbation on body volume cannot inform on mechanisms of size uniformity among individuals. Understanding size uniformity requires precise measurement of individuals at high throughput ^8–11^, which has remained challenging at a multi-cellular scale.

For unicellular systems, a powerful approach to study size control has been the correlative analysis of cell size at birth and division ^12–16^ that distinguished so-called *sizer* and *adder* mechanisms of cell size homeostasis. For a mechanism involving a size threshold, the size at cell division must be independent of a cell’s history, such as the size at cell birth. For *sizers*, the volumes at cell birth and cell division should therefore be uncorrelated. Although evidence for sizers exists for several cell types ^17–19^, most bacteria, yeasts, and cultured mammalian cells do not divide at a fixed size threshold. Instead, many cells functions as *adders*: On average, they add a constant absolute volume in each cell cycle ^12–16,20,21^. Similar to sizers, adders converges to a stable cell size ^12,13,16^.

The adult volume of multi-cellular animals is determined by multiple parameters, including the growth rate (i.e., the rate of volume increase), the duration of development (i.e., the time during which growth occurs before adulthood is reached), and the volume at birth. In principle, each of these parameters can fluctuate independently of the other parameters, and fluctuations in two parameters can either cancel each other out or add up to amplify body volume changes. For example, a slowly growing individual can reach the same volume as a rapidly growing individual if its development is sufficiently slowed down, providing more time for body growth to occur.

In this study, we used *C. elegans* to measure the heterogeneity of volume growth among individuals. *C. elegans* is well-suited for this purpose, given its stereotypic and precisely characterized and development through four larval stages (L1 to L4), and recent technological advances for high-throughput single animal cultivation ^11,22–25^. Moreover, most somatic cells of *C. elegans* are post-mitotic, such that volume growth is predominantly due to increase in cell size, rather than cell proliferation ^26^, which facilitates the formulation of quantitative models of body growth.

The duration of *C. elegans* larval stages is thought to be controlled by a developmental clock with oscillatory transcriptional output ^23,27,28^. Although the molecular mechanism driving these oscillations remains unknown, the output of the developmental clock has been well characterized: the clock controls the oscillatory expression of >3000 genes. All genes share the same oscillatory period of ∼8 hours, matching the larval stage duration ^23,27^. However, different genes peak at different times, depending on when they function during the moulting cycle of a larval stage. For example, the phase-shifted oscillatory expression of different cuticular collagens ensures their timely during synthesis prior to moulting ^27^. Hence, development and oscillations are tightly linked ^23^: conditions that change the duration of development also change the frequency of oscillations, and vice versa ^27^.

Using live imaging of hundreds of individuals, we characterized the sources of body volume heterogeneity. We found that, unlike unicellular systems, the total body volume of *C. elegans* follows neither a *sizer* nor an *adder* mechanism. Instead, the volume fold change within one larval stage was nearly invariant with respect to the volume at the larval stage entry. Despite the lack of size-dependent growth control by *adders* or *sizers*, we observe very little divergence of body volume between rapidly and slowly growing individuals since the growth rate and the duration of development are anti-correlated. We suggest that *C. elegans* maintains body size homeostasis by coupling of growth and development, and not by a mechanism that responds to size deviations *per se*. Using a mathematical model of developmental oscillations, we show that an inherent dependence of the oscillation frequency on the growth rate can explain this coupling of growth and development.

## Results

### Measurement of growth and body size of individual *C. elegans* larvae

To measure the growth rate and the body size of individuals of *C. elegans*, we used agarose-based microchambers ^22,23^ to track individual animals at high temporal resolution throughout post-embryonic development by live imaging. We used a strain ubiquitously expressing *gfp* under control of the *eft-3* promoter from a single-copy transgene ^23^, providing high contrast images for robust and precise segmentation of the outline of animals (Fig. 1b). We placed individual embryos of this strain in arrayed chambers filled with the bacterial strain *E. coli* OP50, the standard laboratory diet of *C. elegans*, and sealed these chambers by adherence to a gas permeable polymer. This experimental setup allowed us to image up to 250 individuals of *C. elegans* in parallel at a time resolution of 10 minutes from birth to adulthood by fluorescence microscopy. The temperature was kept constant at 25°C +/-0.1°C by enclosing the microscope in a dedicated incubator. In total, we collected data of 1153 individuals from 12 micro chamber arrays that were imaged on 10 different days.

We determined the body volume at each time point from 2D images, by computationally straightening the central focal plane of each worm after segmentation and assuming rotational symmetry, as previously described^29^ (Fig. 1b,c). Consistent with previous observations ^5^, we observed four plateaus with near absent or even negative volume growth for ∼2 hours, followed by a saltatory increase in body volume (Fig. 1c). This halt in growth was accompanied by a lethargic period prior to cuticular ecdysis^30,31^, during which animals stopped feeding with a constricted pharynx and intestine ^30^ (Fig. S1a). We could thereby determine larval stage transitions as the timepoints of growth restart after each lethargic episode. The average larval stage durations determined using this assay were close to manual assignments in microchambers ^23^, and on standard petri-dishes ^32^ (mean duration was 13.1 h, 8.0 h, 7.7 h, 10.4 h for L1-L4). On average, animals increased 40-fold in volume between birth and the last moult, with 2-to 3-fold volume changes per larval stage. Volume fold changes were larger in L1 and L4 stages than in L2 and L3, consistent with the longer duration of these two stages (Fig. S1b).

### *C. elegans* grows faster than linearly within larval stages

Body volume uniformity among individuals is especially sensitive to changes in the growth rate during exponential growth, where the absolute volume increase per time is proportional to the current volume. Hence, under exponential growth, the difference in volume between two individuals with different growth rates amplifies exponentially with time (Fig. 1a). Previous research has described growth of *C. elegans* as piecewise linear, with constant linear growth within a larval stage and a saltatory increase in the linear growth rate upon larval stage transitions ^5,33^. However, we note that distinguishing between linear and exponential growth within a larval stage requires highly accurate measurements^34^. Moreover, measurements at high temporal resolution are needed to avoid confounding effects of growth pauses during lethargus.

To address if the body volume of *C. elegans* increases linearly, or faster than linearly within a larval stage, we determined the average absolute rate of volume increase 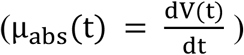 and the rate of volume increase normalized to the current volume 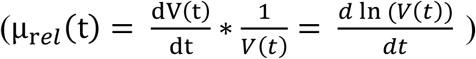 within each larval stage. To this end, we divided the larval stages of each individual into 100 equally spaced intervals and averaged µ_abs_ and µ_rel_ in each interval over all individuals. This analysis yielded the growth rate as a function of larval stage progression τ. Consistent with supra-linear growth, the mean absolute growth rate <µ_abs_(τ)> increased monotonically during development with the exception of the lethargic periods immediately prior to ecdysis (Fig. 1d). Consistently, he average relative growth rate <µ_rel_(τ)> was nearly constant within larval stages L1 to L3, consistent with exponential growth. Growth during the L4 stage was also faster than linear (Fig. 1d), but slower than exponential, as µ_rel_ declined towards the end of the larval stage (Fig. 1d). Notably, this decrease of µ_rel_ coincided with a previously reported slowing down of the oscillatory developmental clock at the end of L4 ^23^. The decline of relative growth rate during L4 is therefore likely a developmental feature of this larval stage, although we cannot entirely exclude an influence of geometric or other constraints on µ_rel_.

To exclude that supra-linear growth within a larval stage was caused by re-scaling and averaging, we fitted a linear and an exponential growth model to each individual. We excluded the first 10% and the last 25% of each larval stage to avoid confounding effects of the growth halt during lethargus and the saltatory volume increase after moulting. For L1 to L3 stages, more than 90% of the individuals had a better fit to the exponential growth model (higher pearson correlation coefficient between fit and data for 98%, 99%, and 93% of individuals of L1-L3, p<10^−50^ bionomial test). For the L4 stage, linear growth provided a better fit than exponential growth for the majority of individuals (83%, p<10^−50^, binomial test), confirming that, although growth is faster than linear during the L4 stage (Fig. 1d), the growth dynamics during L4 cannot be captured by a linear nor by an exponential growth model.

Finally, we found that for all larval stages, individuals with a large body volume at the beginning of the larval stage have a faster absolute rate of volume increase µ_abs_ than smaller animals of the same larval stage (Fig. S1c,d), as is expected for autocatalytic growth. We conclude that *C. elegans* grows faster than linearly during all larval stages, raising the challenge of amplifying heterogeneity in body volume during development due to differences in the growth rate. In the following, we will refer to µ_rel_ as the *growth rate* without specifying normalization to the current volume at every use. µ_abs_ will be specified as the *absolute growth rate*.

### Maintenance of body size uniformity despite growth rate heterogeneity

We next asked how much individuals of *C. elegans* differed among each other in their growth rate. Heterogeneities among individuals can either stem from batch effects of day-to-day repeats, or from heterogeneity among individuals from the same batch. Despite highly standardized experimental conditions, we observed small but significant differences between batches that may stem from microenvironmental, parental ^8^, or other effects. To exclude any influence of batch effects on our conclusions, we normalized growth and size relative to the mean of the population of each respective day. For the rest of this article, main figures will show results from batch corrected data, whereas the related Supplemental Figures will show that the trends hold even without batch correction, in the unnormalized data.

After normalization, the coefficient of variation (CV) in the growth rate ranged between 6% and 12%, depending on the larval stage (Figs. 2ab, 4d). For a given individual, the differences in the growth rates persisted between consecutive larval stages (Fig. S2a). Autocorrelation was reduced when comparing larval stages further apart and was nearly absent when comparing L1 with L4 larvae. Hence, genetically identical individuals of *C. elegans* display heterogeneity in their growth rate that is partially transmitted between larval stages with decay of the autocorrelation on a time scale of days. We observed a similar correlation for the duration of the larval stages across development (Fig. S2b), consistent with previous observations ^32^. Positive autocorrelation of growth rates also indicates that *C. elegans* does not undergo catch-up growth^7^, where individuals that grow slowly in the beginning of development catch up by growing faster later in development, although we cannot entirely exclude that micro environmental drive positive auto-correlation.

**Figure 2.**
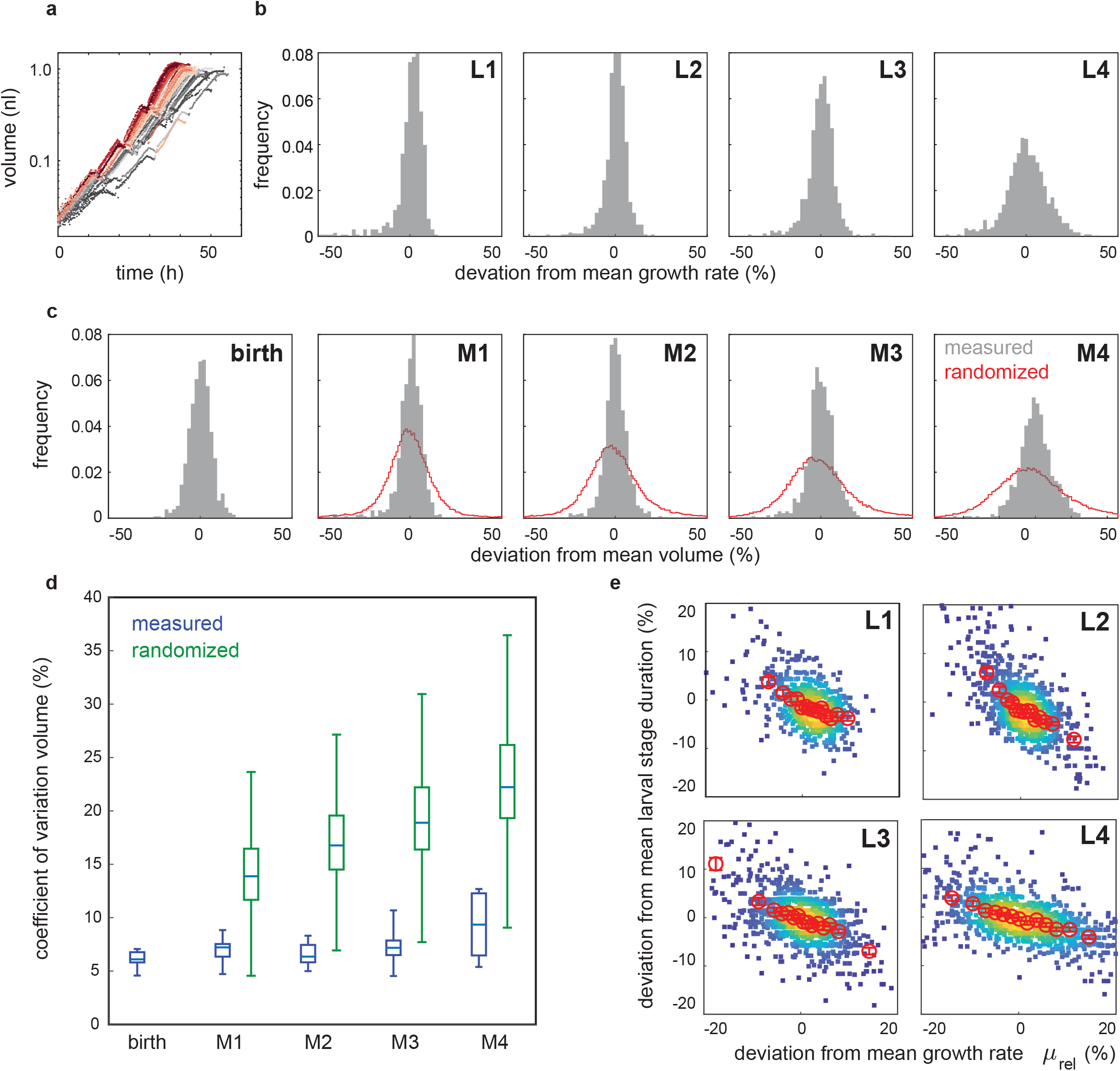
Volume divergence among individuals is less than expected by the heterogeneity in growth rates. *(related to Supplemental Figure S2)* a. Example of volume trajectories for all individuals measured of one experimental repeat. Colours represent the rank of growth rate at each larval stage (grey = slowest, red = fastest). c. Histogram of the growth rate deviations in % from the population mean measured at indicated larval stages. The growth rate of each individual and larval stage was determined by a linear regression to log(volume) against time excluding the first 10% and the last 25% of the larval stage. c. Histogram of body volume deviations in % from the mean measured at birth and indicated larval moults. Red line shows distribution of volume resulting from random shuffling of growth rate and larval stage duration among individuals. d. Coefficient of variation of volume during development. Blue: box plot of the average CV measured experimentally in the different day-to-day repeats. Green: expected CV based on 1000x random shuffling of growth rate and larval stage duration. The measured CV was significantly smaller than the randomized control at all stages (p<10^−5^, ranksum test) e. Scatter plot of growth rate vs. larval stage duration shown as % deviation from the mean. Colour indicates point density. Red circles are moving average along x-axis +/- SEM.

To ask if differences in the growth rate accumulate to differences in volume during development, we next determined the heterogeneity in body volume at each larval stage. At birth, the CV of the body volume was 6.1% and increased moderately to 9.6% at L4-to-adult transition (Figures 2c,d, S2d,e). This increase in body volume heterogeneity was significantly smaller (p<10^−5^, ranksum test) than expected based on random shuffling of growth rates and larval stage durations (Fig. 2c,d, S2d,e, see methods). Our measurements therefore suggest a mechanism that buffers body volume against heterogeneities in the growth rate. Indeed, the duration of larval stages was anti-correlated with the growth rate of individual animals (Fig. 2e, R^2^=0.51, 0.56, 0.43, 0.41 for L1 to L4, p<10^-5^ for all stages), consistent with previous measurements of a small number of individuals in liquid growth medium ^5^.

### Invariance of the volume fold change within larval stages

Size homeostasis has often been associated with size thresholds for passing developmental milestones^4–7^. Building on previous analysis of unicellular systems^12–17,19–21^, we investigated the importance of size thresholds during *C. elegans* development by analysing the relation between the volume at the beginning (V_1_) and the volume at the end (V_2_) of each larval stage. If larval stage progression was determined by a fixed size threshold V_T_ then V_2_ should be history-independent, such that V_2_ and V_1_ are uncorrelated. For an *adder* mechanism, V_2_ correlates positively with V_1_ and the absolute volume increase (ΔV = V_2_-V_1_) per a larval stage is independent of V_1_ (Fig. S3). To distinguish adders and sizer during *C. elegans* growth, we analysed the correlation between relative deviation of V_1_ and V_2_ from the population mean for each individual. This normalization allowed us to quantitatively compare the relation between V_1_ and V_2_ among different larval stages despite the large volume differences between L1 and L4 larvae.

We found two distinct modes of volume control, depending on the larval stage. The L1 stage behaved close to an adder, with ΔV being nearly independent of V_1_ (Fig 3b). However, for L2 to L4 stages, the data matched neither an adder nor a sizer mechanism since V_2_ and ΔV were both positively correlated with V_1_ (Fig. 3a-b, Fig. S4). Instead, for these stages, the volume fold change (FC_V_ = V_2_/ V_1_) was nearly independent of V_1_ (Fig. 3c, Fig. S4) with the exception of the smallest L2 larvae (Fig 3c). In analogy to adders and sizers, we term this relation a *folder* mechanism.

**Figure 3.**
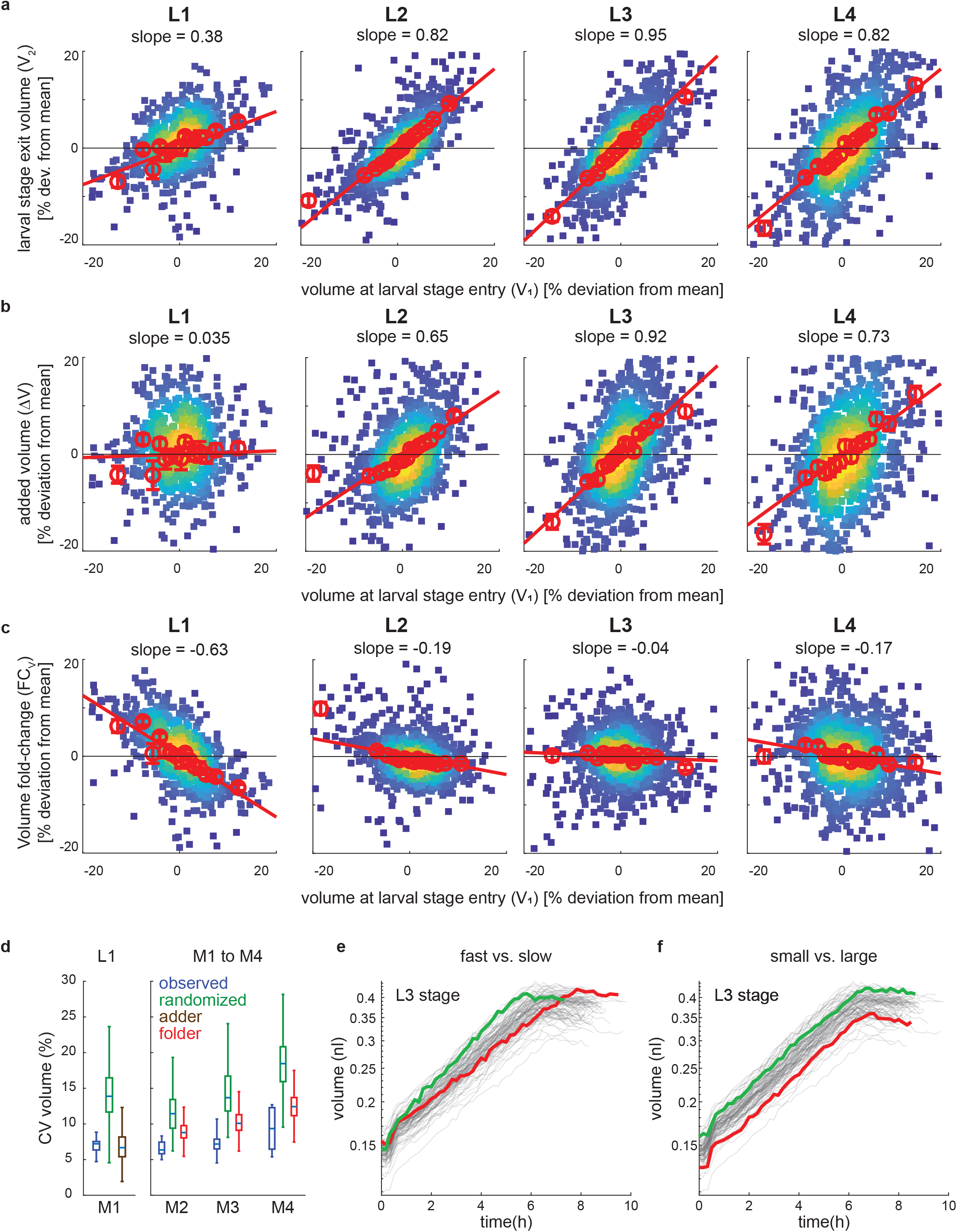
Constant volume fold change per larval stage for L2 to L4. *(related to Supplemental Figure S3 and S4)* a. Scatter plot of volume at larval stage entry vs. volume at larval stage exit shown as % deviation from the mean for indicated larval stages. Colour indicates point density. Red circles are a moving average along x-Axis +/- SEM. Red trendline: a linear regression to moving average, excluding the first and the last point. Slope of the trendline is indicated above the figure. b. Same as (a), but for volume at larval stage entry vs. absolute volume increase per larval stage. c. Same as (a), but for volume at larval stage entry vs. volume fold change per larval stage. d. Comparison of CV of volume to adder and folder models. Blue: box plot of the average CV measured experimentally in the different day-to-day repeats. Green: expected CV based on 1000x random shuffling of growth rate and larval stage duration. Brown: CV expected from adder model during L1. Red: CV expected from folder model during L2 to L4. Randomizations was done separately for L1 and for L2 to L4 starting from measured volume distributions at birth and at M1, respectively. e. Illustration of folder phenomenon. Grey lines: volume as a function of time of the L3 stage of all individuals of one experimental repeat. Two highlighted individuals (red and green) with similar starting volume, but different growth rates reach the same volume at the end of the larval stage. f. same as (e), but highlighting two individuals with similar growth rates, but distinct starting volumes that maintain the same relative volume difference at the start and end of the larval stage.

### Two rules describe growth of *C. elegans*

Thus, our temporally highly resolved growth measurements of individual animals have revealed two fundamental rules of *C. elegans* growth. Rule #1: Larvae undergo a volume fold change that is independent of their volume V_1_ at larval stage entry. Rule #2: Larvae exhibit little increase of body volume heterogeneity across individuals over the course of development despite significant heterogeneity in their growth rates. In principle, any exponential growth for a fixed amount of time t could explain rule #1. More generally, volume fold changes that are independent of could even occur for uncorrelated µ and t 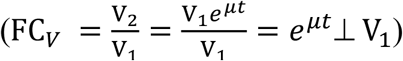 However, for a fixed timer or for uncorrelated µand t, the differences in volume between individuals with distinct growth rates are expected to increase during development, which would be inconsistent with rule #2.

The observed uniformity in body volume (rule #2) is not due to size-dependent regulation of growth, as we find no evidence that small individuals catch up in size within a larval stage (Fig. 3a,b). Instead, volume divergence is prevented by an anti-correlation of the growth rate and the larval stage duration t (Fig. 2d). Hence, unlike adders and sizers, a folder does not compensate for differences in size *per se*, but merely prevents the exponential divergence in volume between slowly or rapidly growing individuals. Figures 3e and 3f highlight stereotypic examples of two L3 stage individuals differing in either their growth rate µ (Fig. 3e), or in their initial volume V_1_ (Fig. 3f) to illustrate the implications of the folder mechanism on body size divergence.

Notably, although a folder slows down divergence in body volume among individuals, in the presence of noise around the average volume fold change, a folder does not entirely prevent volume divergence (Fig. S3d). Intuitively, this imperfection can be understood by the absence of size dependent control, such that stochastic deviations from the appropriate volume are propagated to later stages. Thus, the behaviour of a folder is distinct from adders and sizers observed in cells, which converge to a stable volume over multiple cell cycles even if there is noise in V_T_ or ΔV ^12–17,19–21^.

To estimate the divergence in volume expected from a combined adder and folder mechanism, we simulated volume divergence by the adder during L1 and by the folder during L2 to L4 by randomly shuffling ΔV (for L1) and FC_V_ (for L2 to L4) among individuals. As expected, the simulated volume divergence by a folder was substantially smaller than completely random shuffling of growth rates and developmental times and close to experimental observations (Figs. 2d, S2e, red). The experimentally observed volume divergence was even smaller than the simulated folder (Fig. 3d), which may be explained by a weak correlation between FC_v_ and V_1_ (Fig. 3c) or by more complex processes.

### Coupling of growth and development is robust to changes in growth rates

Since the folder mechanism relies on a coupling between the rates of growth and development, we asked if body size homeostasis is impaired by genetic manipulation of known pathways of growth control and developmental timing. First, we used a mutation of *eat-2*, which impairs growth by a reduction of pharyngeal pumping, and thus of food intake ^35^. Second, we impaired mTOR signalling by a deletion of the RagA homolog *raga-1* ^36,37^. Third, we perturbed TGFβ signalling by overexpressing the TGFβ ligand DBL-1 ^38^ and by a mutation of the TGFβ target *lon-1*^3,39^. Fourth, we perturbed developmental timing by a mutation in the heterochronic gene *lin-14* ^40^.

To compare different mutants with each other, we define the rate of development α as the inverse of the larval stage duration t (α = 1/t). Mutations with proportional effects on α and on the growth rate µ do not alter the volume fold change 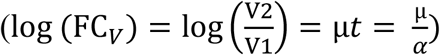. Differences in the body volume among mutants can therefore stem from non-proportional changes to μ and α, or from differences in the volume at birth.

Mutants differed only minimally from the wild type in their volume at birth (Fig. 4b) but differed substantially among each other and from the wild type in their rates of growth and development (Fig. 4a). Overall, mutant effects on μ and α were strongly correlated (Fig. 4a, R = 0.91, 0.89, 0.63, 0.95 for L1 to L4), indicating a coupling of growth and development across different genotypes. However, individual mutants deviated from perfect proportionality between α and μ, resulting in significant alterations in body volume (Fig. 4b). For example, mutation of *eat-2* consistently slowed down the growth rate more strongly than the rate of development (Fig. 4a), resulting in animals that were smaller than wild-type animals of the same larval stage (Fig. 4b). *lon-1* mutants were near proportionally affected in growth and development and did not change in final volume (Fig. 4b), although they were significantly longer and thinner than wild-type animals (Fig. 4c)^41^. Finally, *lin-14* mutation reduced μ less strongly than α, such that *lin-14* mutants were larger than wild-type animals after three larval stages. Since *lin-14* mutant animals undergo only three larval stages ^40^, *lin-14* mutants were nevertheless smaller than wild type at transition to adulthood (Fig. 4b). Together, these data show that genetic mutations can affect the rate of growth and the rate of development non-proportionally, which alters the adult body volume.

**Figure 4.**
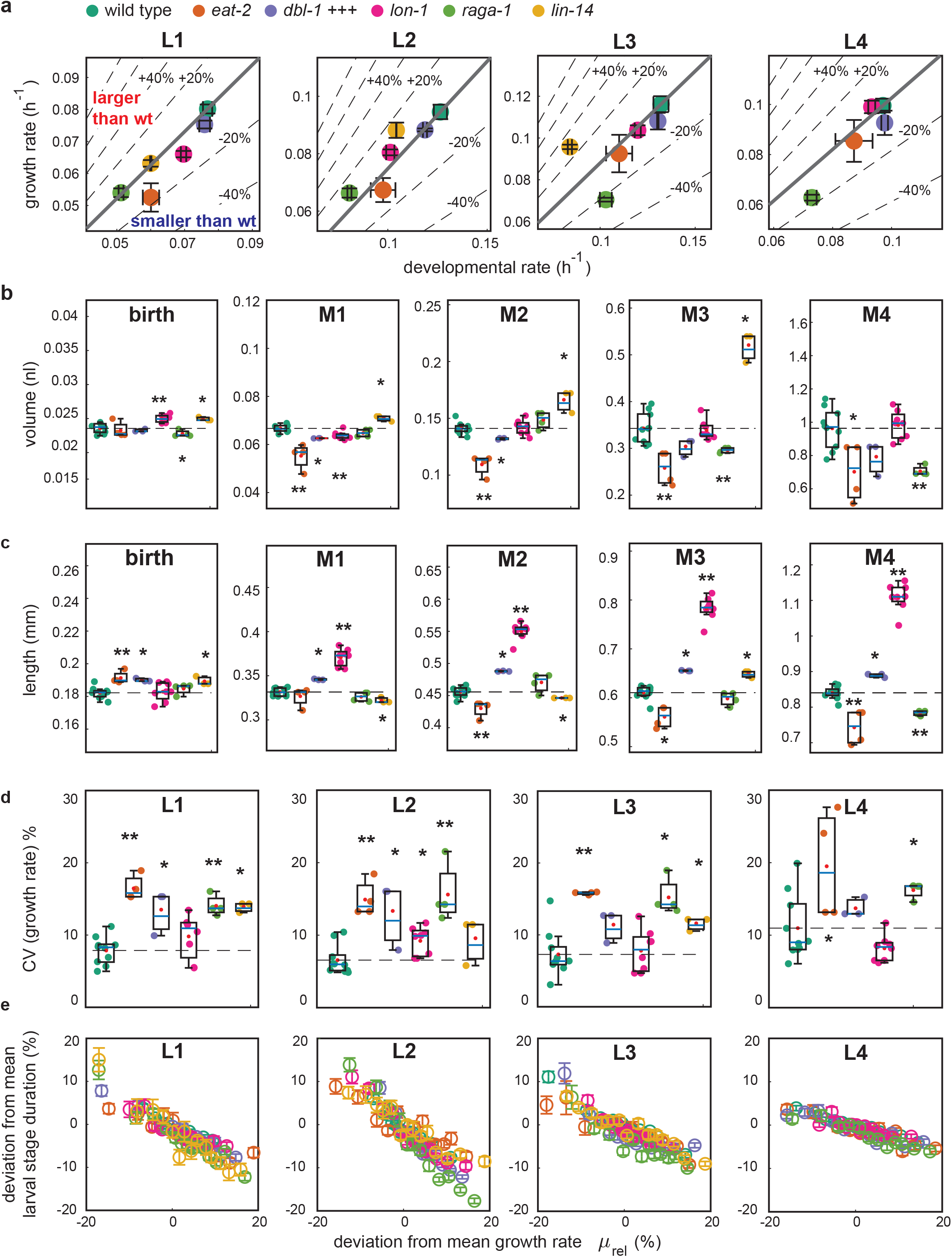
Mutations that alter growth and body size do not uncouple growth and development. *(related to Supplemental Figures S4 and S5)* a. Scatter plot of the rate of development (=1/larval stage duration) vs. the growth rate for indicated mutants and larval stages. Colour scheme as indicated in legend of (b). Error bars: SEM between day-to-day repeats. Thick grey line indicates proportional scaling corresponding to the volume fold change equivalent to the wild type. Dotted lines indicate deviation in volume fold change for a given deviation from proportionality. Region above the thick line corresponds to an increase in volume fold change, region below the thick line to a decrease in volume fold change compared to the wild type. b. Box plot of volumes at birth and larval moults for indicated mutant strains. Individual points and box plots indicate average volume of each day-to-day repeat. Significance is calculated by ranksum test of day-to-day repeats between mutant and wild type (* = p<0.05, ** = p<0.005). c. same as (b), but for body length. d. same as (a), but for coefficient of variation of the growth rate. e. Correlation between growth rate and the larval stage duration among individuals for different mutant backgrounds shown as deviation from the mean. Colours correspond to legend shown in (b). Circles show moving average along the x-Axis +/-SEM, as described in Figure 3. Scatter plot of individuals is omitted for clarity of presentation (see Fig. S5).

We next asked if a non-proportional change of α and μ would also disrupt the anti-correlation of growth and development among different individuals of the same genotype. However, for all mutant strains, the growth rate and the duration of larval stages remained anti-correlated with a quantitative relationship close to that of wild type animals (Fig. 4e, Fig. S5). Although several mutants had strongly impaired uniformity in their growth rate (Fig. 4d) the volume divergence of mutants remained smaller than expected by chance and was even slightly smaller than expected by the folder model (Fig. S6). One exception to this rule was the *eat-2* mutant, for which divergence was close to expectations from random shuffling from the L2 stage onwards (Fig. S6).

In summary, we conclude that the coupling of growth and development among individuals is robust to impairment of growth rate uniformity and occurs independently of the pathways investigated (Fig 4e, Fig. S5). Quantitative differences in the volume divergence among mutants may relate to heterogeneities in traits that cannot be determined by volume measurements alone.

### The folder mechanism temporally coincides with the onset of developmental oscillations

Since perturbation of canonical growth control pathways did not impair the coupling of growth and development, we speculated that such coupling could instead be mediated through the oscillatory clock of *C. elegans* development^23,27^. To test this hypothesis, we focused on the L1 stage which, unlike other larval stages, follows an adder and not a folder mechanism (Fig. 3). The L1 stage also differs from other stages with respect to the developmental oscillator: During L2 to L4 stages, the oscillations are in synchrony with larval stages, whereas the oscillator is arrested during the first 5 to 7 hours of the L1 stage ^23^ (Fig. 5a). We therefore asked whether transition to the folder mechanism temporally coincided with the onset of gene expression oscillations.

**Figure 5.**
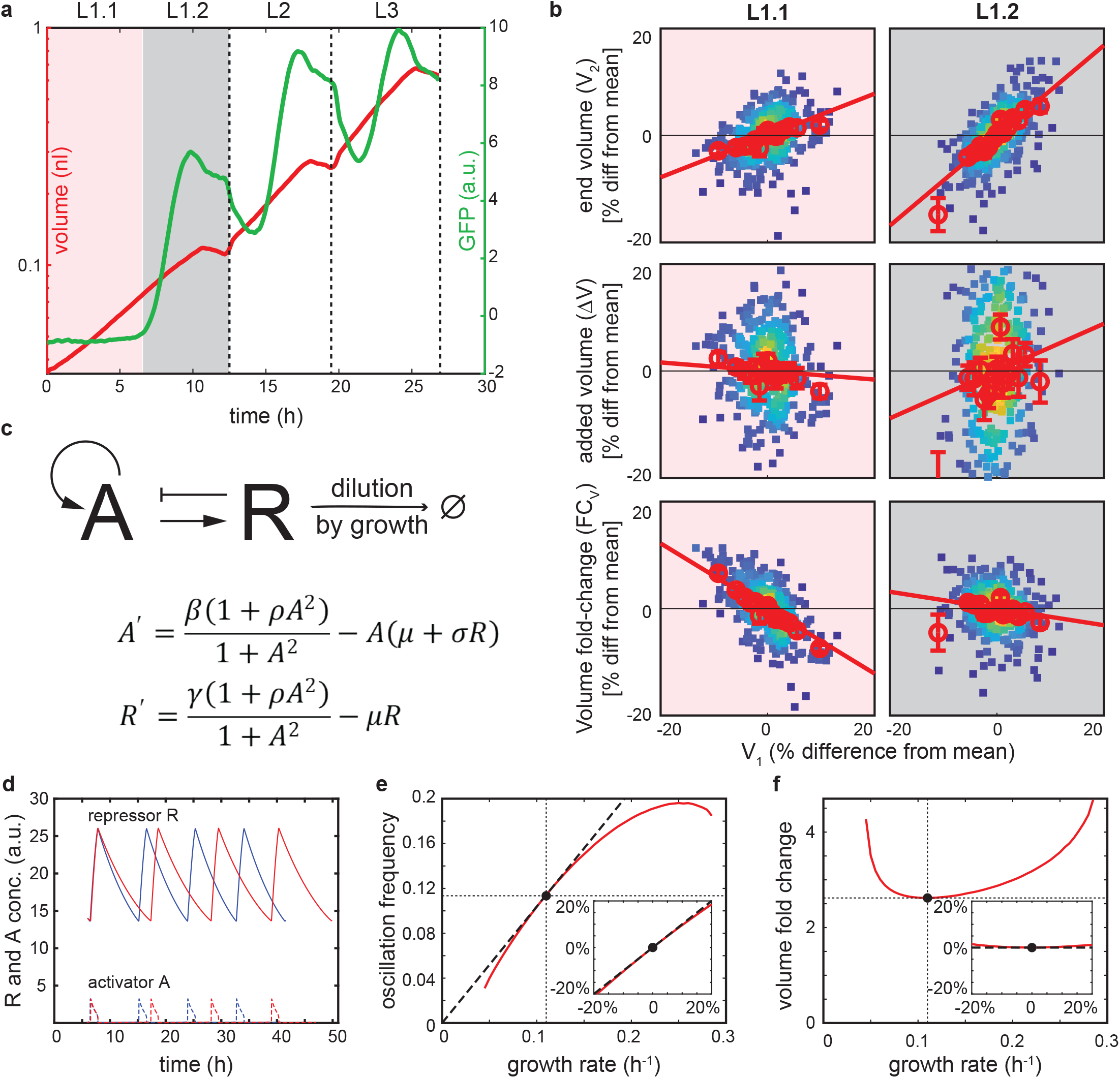
Developmental oscillations temporally coincide with and explain the folder. *(related to Supplemental Figure S6)* a. Measurement of GFP concentration (green) and volume (red) of *dpy-9*:*GFP* strain for larval stages L1 to L3. L4 stage was not measured in this experiment due to technical constraints. Figure shows GFP concentration and volume averaged over all individuals after temporal re-scaling of each stage separately to align individuals with slightly different growth rates. After re-scaling and averaging, the data was scaled back to the average larval stage duration. L1 is split into pre-oscillation (red, L1.1) and post-oscillation (grey, L1.2) substages. b. Adder and sizer comparison for L1.1 and L1.2 as described in Figure 3. c. Mathematical model of genetic oscillator based on design II of ref. ^43^. Activator *A* activates its own production and the production of the repressor *R* by a factor of ρ. *R* degrades *A* at a rate σ. β and γ are the basal production rates of *A* and *R*, respectively. µ is the growth rate. *R* is considered stable, such that its removal rate of is set by the growth rate µ. Model parameters (β = 10/h, γ = 0.3/h, σ = 10/h, ρ = 50) are close to a saddle-node on invariant circle (SNIC) bifurcation ^45^ as is experimentally observed ^23^. The invariance of the volume fold change with respect to the growth rate is qualitatively independent on parameters over a wide range (see Fig. S6). d. Modelled dynamics of *A* (dotted lines) and *R* (sold lines) for µ = 0.12/h (blue), and for µ = 0.10/h (red). e. Oscillation frequency as a function of the growth rate (red). Missing values at slow and fast growth rates are because the system is not oscillating at these extremes. Black dotted line: perfect proportionality between the growth rate and the oscillation frequency. The black circle: reference growth rate at which proportional scaling is precise for the given parameters (µ =0.11/h). Insert shows the same data but mean normalized to the reference growth rate +/- 20%, equivalent to the display of experimental observations in (b) and Figure 3. f. Same as (e), but for growth rate vs. volume fold change.

To this end, we imaged growth of a strain expressing *gfp* under control of the oscillatory collagen promoter *dpy-9p* ^23^ (Fig. 5a). As expected, *dpy-9p*::*gfp* expression was undetectable during the first 6 hours of the L1 stage when the oscillator is arrested ^23^, and *gfp* expression oscillated in synchrony with larval stages later in development (Fig. 5a). Using *dpy-9p*::*gfp* expression, we divided the L1 stage into two substages (L1.1 and L1.2) before and after oscillations start. L1.1 behaved close to an adder (Fig. 5b), similar to what we observed for the entire L1 stage (Fig. 3b). In contrast, L1.2 was close to a folder (Fig. 5b), similar to stages L2 to L4 (Fig. 3c). Hence, the folder mechanism occurs specifically during the developmental window of active oscillations, supporting a role of developmental oscillations in the coupling of growth and development.

Although the molecules that control developmental oscillations in worms are unknown, many biological oscillators are driven by an ultrasensitive feedback between an activator protein A and a repressor protein R (Fig. 5c) ^42^. The frequency of such oscillators is strongly dependent on the removal rate of the repressor R ^43^. Since, for stable proteins, the removal rate is equivalent to the rate of dilution through growth ^44^ the frequency of oscillations may be directly influenced by the growth rate if R is stable (Fig 5d). To test this, we analysed a model of a biological oscillator (Fig. 5c, based on ^43^) operating close to a saddle-node on invariant circle (SNIC) bifurcation ^45^, as has been observed experimentally ^23^. In this model, the oscillation frequency indeed scaled near proportionally with the growth rate over a wide range (Fig. 5e), producing a near constant volume fold change (Fig. 5f) independent of the model parametrization (Fig. S7).

In conclusion, we propose a minimal model where the coupling of growth and development emerges as an intrinsic property of a developmental oscillator that minimizes body volume divergence among individuals with different growth rates. Identifying the molecular components that drive developmental oscillations will enable testing of this model and may allow for predictive modulation of organismal body volume.

## Discussion

Exponential growth of cells and organisms presents the challenge that small differences in the growth rate can, in principle, amplify to large differences in size. Cells overcome this challenge by adder and sizer mechanisms, where the volume fold change per cell cycle correlates negatively with the size at birth ^12–17,20,21^. Here, we found that, for *C. elegans*, size uniformity at an organismal scale follows a different principle. *C. elegans* neither maintain a fixed volume added per larval stage (adder) nor a fixed size threshold (sizer) at which moulting occurs. Instead, individuals delay or accelerate moulting depending on their growth rate, such that, e.g., rapidly growing individuals grow for a shorter amount of time and vice versa. This behaviour results in a volume fold change per larval stage that is invariant with respect to the starting size (Fig. 3). Despite the apparent lack of size dependent growth control, we observed only a minor divergence of body volume between rapidly and slowly growing individuals (Fig. 2d), suggesting the coupling of growth and development as a potent mechanism for body size homeostasis.

Our observation of coupling of growth and development is consistent with the recent observation that the temporal spacing of different morphologically defined events of *C. elegans* development scale proportionally among individuals ^46^. Our model further suggests that the time between developmental events is determined by the rate of exponential growth of an individual, and that differences in the growth rates among individuals explain the corresponding differences in the timing of developmental events. We thereby propose the growth rate as a central regulator of organismal physiology and development, similar to growth laws found in bacteria ^47,48^.

How could animals achieve coupling between growth and development? At least two distinct, but not mutually exclusive, mechanistic models are possible. First, mechanical stretching of structural components, such as the cuticle, could trigger moulting events in ways that maintain a constant fold change ^49^. Alternatively, the growth rate may influence the dynamics of a developmental clock, such as the developmental oscillator of *C. elegans* ^23,27^. In support of a role of the oscillator in the folder mechanisms, we showed that the folder is restricted to the developmental window where the oscillator is active (Fig. 5a,b). Moreover, we show that in a minimal mathematical model of a genetic oscillator the oscillation frequency scales near proportionally with the growth rate if the degradation rates of the oscillator components are small (Fig. 5e). Thus, a circuit design with only two components and without complex regulatory architecture is sufficient to explain the coupling of growth and development we observed. We of course do not exclude that additional, more complex regulatory mechanisms may be at play. Nevertheless, this minimal model makes quantitative predictions that will help to identify the molecules driving transcriptional oscillations in *C. elegans*. For example, we predict that increasing the degradation rate of the repressor R increases oscillatory frequencies, thereby shortens development, and hence reduces body size. In contrast, perturbations of growth that are unrelated to the oscillator are predicted to have a much smaller effect on the body size.

Transcriptional oscillations are found in numerous organisms. Most famously, the circadian clocks controls oscillations that match the diurnal cycle ^50^. Unlike the developmental clock of *C. elegans*, the 24 hour period of circadian clocks is highly robust and independent of fluctuations in growth rates or temperature ^50^. We propose that, for developmental oscillations, the apparent lack of robustness to changes in growth rates serves the purpose of ensuring robustness of body size homeostasis. It will be interesting to see if this fundamental design principle also applies to the size homeostasis of other multicellular systems controlled by oscillations, such as somite formation in vertebrates ^51^.

## Experimental procedures

### Caenorhabditis elegans strains

#### The following strains were used in this study

HW1939: *xeSi296[eft-3p::luc::gfp::unc-54 3′UTR, unc-119(+)] II* (^23^)

HW2688: *xeSi296[eft-3p::luc::gfp::unc-54 3′UTR, unc-119(+)] II; lon-1(e185) III*. (this study)

HW2688: *xeSi296[eft-3p::luc::gfp::unc-54 3′UTR, unc-119(+)] II; lon-1(e185) III*. (this study)

HW2696 *xeSi301[eft-3p::luc::gfp::unc-54 3′UTR, unc-119(+)] III*.; *raga-1(ok386) II*. (this study)

HW2687: *xeSi296[eft-3p::luc::gfp::unc-54 3′UTR, unc-119(+)] II; lon-1(e185) III. ctIs40[[dbl-1(+) sur-5::GFP] X*. (this study)

HW2681: *eat-2(ad1113) xeSi296 [eft-3p::luc::gfp::unc-54 3 ‘UTR, unc-119(+)] II*. (this study)

HW1973: *xeSi296 [eft-3p::luc::gfp::unc-54 3 ‘UTR, unc-119(+)] II*.; *lin-14 (n179)*

HW2840: *xeSi449[eft-3p:mCherry-luciferase unc-119(+)] III*.; *xeSi440[dpy-9p::GFP::H2B::Pest:: unc-54 3 ‘UTR; unc-119(+)] II*. (this study)

The GFP reporters *xeSi301* is equivalent to *xeSi296* ^23^, except that it was inserted on chromosome III instead of chromosome II. *xeSi449* is the same reporter as *xeSi301*, with mCherry instead of GFP. All transgenes were inserted by MosSCI as described in ^52^.

### Microchamber imaging

Imaging of individual animals was performed using a protocol adapted from ^22^ and as described in detail in ^23^ except that a 3.5cm dish with optical quality gas-permeable polymer (ibidi) was used to mount the chambers. In brief, arrayed micro chambers were produced from a 4.5% Agarose gel in S-basal using a PDMS template as a micro comb to create the chambers. Dimensions of chambers used in all Figures except Figure 5 were 600 x 600 x 20 μm. Chambers used in Figure 5 were 370 x 370 x 15 μm for compatibility with camera chip size of the microscope used. Chambers were filled with bacteria of the strain OP-50, which was scraped off a standard NGM plate using a piece of agar supported by a glass slied. After placing individual eggs into the chambers, the micro chamber arrays were inverted onto an ibidi dish for imaging. The remaining space around the dish was covered with 3% low melting temperature agarose dissolved in S-basal, cooled down to 42C prior to application to the dish. The dish was then sealed with parafilm to prevent humidity loss during imaging.

For all experiments except those in Figure 5, a 10x objective was used on an Olympus IX70 wide-field microscope with halogen lamp illumination and an sCMOS camera with a pixel size of 6.5 μm and 2×2 binning. At each timepoint a z-stack of 7 planes with 5 μm spacing was acquired using a piezo-controlled stage. The focal plane with best contrast was automatically selected for further image analysis. Experiments in Figure 5 were conducted on a Yokogawa spinning disc microscope equipped with two EM-CCD cameras and a beam-splitter. GFP and mCherry signals were acquired sequentially with a total time delay of 35ms. This delay was sufficiently short to overlay the two channels without substantial movement of the animal. At each timepoint, a z-stack of 35 μm with 5μm z-spacing was acquired. The volume at each timepoint was computed from the central focal plane. The fluorescence was computed from the sum of all planes divided by the total number of pixels. For each experiment, control animals without a GFP reporter (HW2840) were imaged and fluorescence of developmental stage-matched animals was used to subtract background and autofluorescence of bacteria and worms. For all experiments, the temperature was maintained at 25C using an incubator encapsulating the entire microscope (life imaging services).

### Image analysis

A custom Matlab script was used to segment worms from raw images. The ImageJ “straightening function” embedded in a KNIME workflow was used for straightening. For segmentation, edge detection by Sobel algorithm using the *edge()* function of Matlab was used, followed by connecting endpoints closest to each other to close gaps in the detected outline. After straightening, each image was classified as either an egg, or a worm using a random forest classifier that trained on a small subset of of manually assigned images. This classification also identified cases where straightening failed (e.g. in the case of self-touching animals), which were removed from further analysis.

### Computation of volume and detection of moults

At each timepoint, the volume was computed from straightened images assuming rotational symmetry. The assumption of rotational symmetry is well justified, based on previous measurements ^11^. Timepoints of larval stage transition were determined by the maximum of the second derivative of the logarithm of the volume. Each volume trace was subsequently inspected and curated manually using a custom-made graphical user interface written in Matlab. To minimize measurement errors, the larval volume at each moult was computed by a linear regression of the volume from ten timepoints preceding the moult (for M1 to M4) or regression to 10 timepoints after hatching for the volume at birth.

### Computation of growth rates

To calculate the continuous linear and absolute growth rates shown in Figure 1d, volumes were median filtered with a window of 3 time points and smoothed over 15 timepoints using the *smooth* function with *rlowess* option of Matlab. Individuals were then re-scaled to the duration of the larval stage and averaged after linear interpolation of the signal at 100 points per larval stage.

To calculate average growth rates per larval stage (Fig. 1e) a linear regression of time vs. log(volume) was performed including all timepoints of a larval stage except the first 10% and the last 25% of each larval stage to avoid confounding effects of lethargus. Absolute growth rates were determined by the same procedure, but by a regression to the non log transformed volume.

### Normalization of day-to-day repeats

Where shown as % deviation from the mean, growth rates, volumes, and times were normalized to the mean of each day. The coefficient of variation was computed as the standard deviation divided by the mean of the normalized data.

### Simulation of randomized populations

To determine the expected volume divergence in the absence of coupling of growth and development given the observed heterogeneity in growth and larval stage duration, simulations were carried out with a starting population of the measured body volumes at start. Each individual was then assigned a growth rate randomly drawn from the measured growth rates (Δln(V)/Δt) and a larval stage duration Δt randomly drawn from the measured larval stage durations. From these parameters, the volume at the following larval stage was computed and the process was repeated iteratively until the end of L4. For fair comparison with measured data given effects of day-to-day variation, randomizations were performed for each day-to-day repeat separately. For each simulation, the number of simulated individuals was equal to the number of individuals in the measured data, and randomization was performed 1000 times for each day-to-day repeat. Box plots in Figures 2d, 3d, S2e, and S6 show the distribution of the coefficients of variation of all simulations. An equivalent procedure was applied to simulate the combined adder/folder model, but instead, individuals were assigned ΔV (for L1) and (for L2 to L4) randomly drawn from the measured data. To compute the randomized and folde model data for M1 to M4 in Figures 3d and S6, the starting volume of the simulation were the measured volumes at M1.

### Computation of trendline in correlation between measured variables

To determine the trendline between two measured variables (red line in Fig. 3a-c and Fig. 5, Fig. S3), data was divided into 15 equally sized bins along variable on the horizontal axis, and a linear regression was performed to the mean values of the bins. The two outermost bins were excluded from this fit, as these were often dominated by extreme outliers. Display of all scatter plots is restricted to the region from -20% to +20% for clarity of display. Few individuals were out of this range as apparent in Figure S4.

### Mathematical model of genetic oscillator

To model oscillations, we expanded on a previously published model by Guantos and Poyatos ^43^. The model describes the protein dynamics of an activator A and a repressor R. A activates its production, as well as the production of R by a sigmoidal input function with a Hill coefficient of 2. R accelerates degradation of A by a factor σ. The model describes protein concentrations of A and R assuming quasi steady-state for the respective mRNAs. In addition to active degradation of A by R, A and R are diluted by growth at a rate µ. β and γ are the basal production rates of A and R, and ρ is the factor by which A enhances the production of its target. Parameters of the model were: β = 10/s, γ = 0.3/s, σ = 10., ρ = 50. Conclusions were qualitatively robust to changes in these parameter values.

To calculate the oscillation frequency a, the dynamics of A and R were numerically solved using Matlab for the range of values of µ where the system adopted limit cycle oscillations. Volume fold changes were computed as FC_v_ = e^µt^ = e^µ/a^.

## Acknowledgements

We thank Iskra Katic for help in generating transgenic strains; Laurent Gelman, Stephen Bourke, and Jan Eglinger for help with imaging and image analysis; Gregory Roth for helpful discussions. B.D.T is a recipient of a HFSP LTF (000309/2013), Marie-Curie IF (#751878), and a fellowship by the Engelhorn-Traudl Foundation. This work received funding from the Swiss National Science Foundation (SNSF) in the form of an Eccellenza Professorial Fellowship (PCEFP3_181204) to B.D.T., from the Novartis Foundation for medical-biological Research (Grant #20A011) and from the European Research Council (ERC) under the European Union’s Horizon 2020 research and innovation program (Grant agreement No. 741269, to H.G.). The FMI is core-funded by the Novartis Research Foundation.

## Author contributions

B.D.T. performed all experiments, data analysis, and modelling and wrote the manuscript. H.G. and

B.D.T. jointly conceived the project and H.G. edited the manuscript.

## Supplemental Figure legends

**Figure S1.**
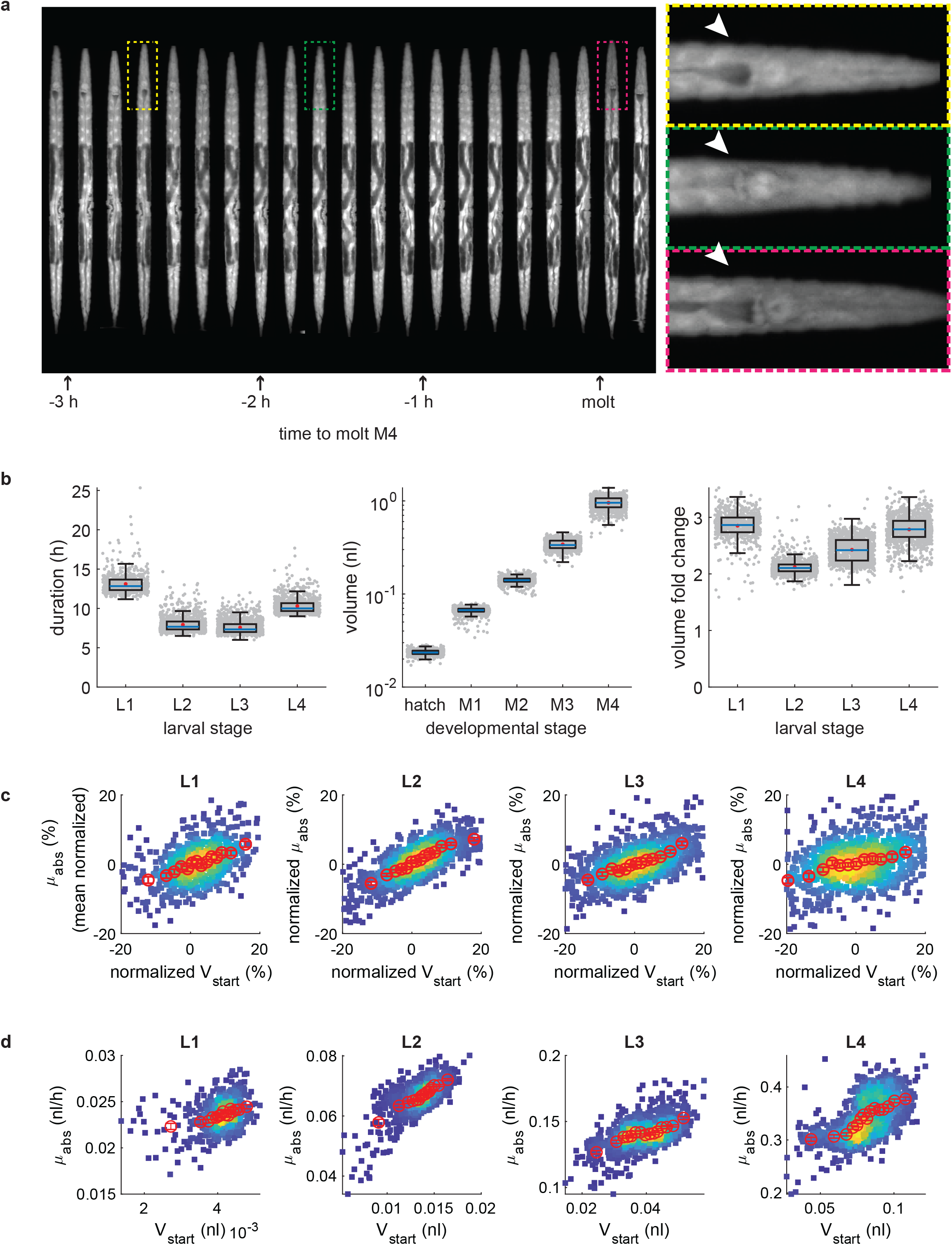
Quantification of volume growth of *C. elegans*. *(relates to Figure 1)* a. GFP signal of a representative time series of straightened images of an individual starting 3 hours before the M4 moult. Yellow box: last time point before lethargus. Green box: time point during lethargus. Pink box: First time point after lethargus. Arrowheads highlight an opening of the intestine posterior to the pharynx (dark area in yellow and pink box) that is not visible during lethargus (green box) when the intestine is constricted and feeding stops. Dark areas in the center of the worm correspond to the gonadal arms, where the *eft-3*::GFP transgene is silenced. b. Box plots of Larval stage duration, volume, and volume fold change in micro chambers. Grey circle are individuals, blue line: median, red circle: mean. c. Scatter plot of absolute growth rate vs. the volume at larval stage entry shown as deviation from population mean. Red circles are the moving average along the x-Axis. c. As (c), but with unnormalized volumes and growth rates.

**Figure S2.**
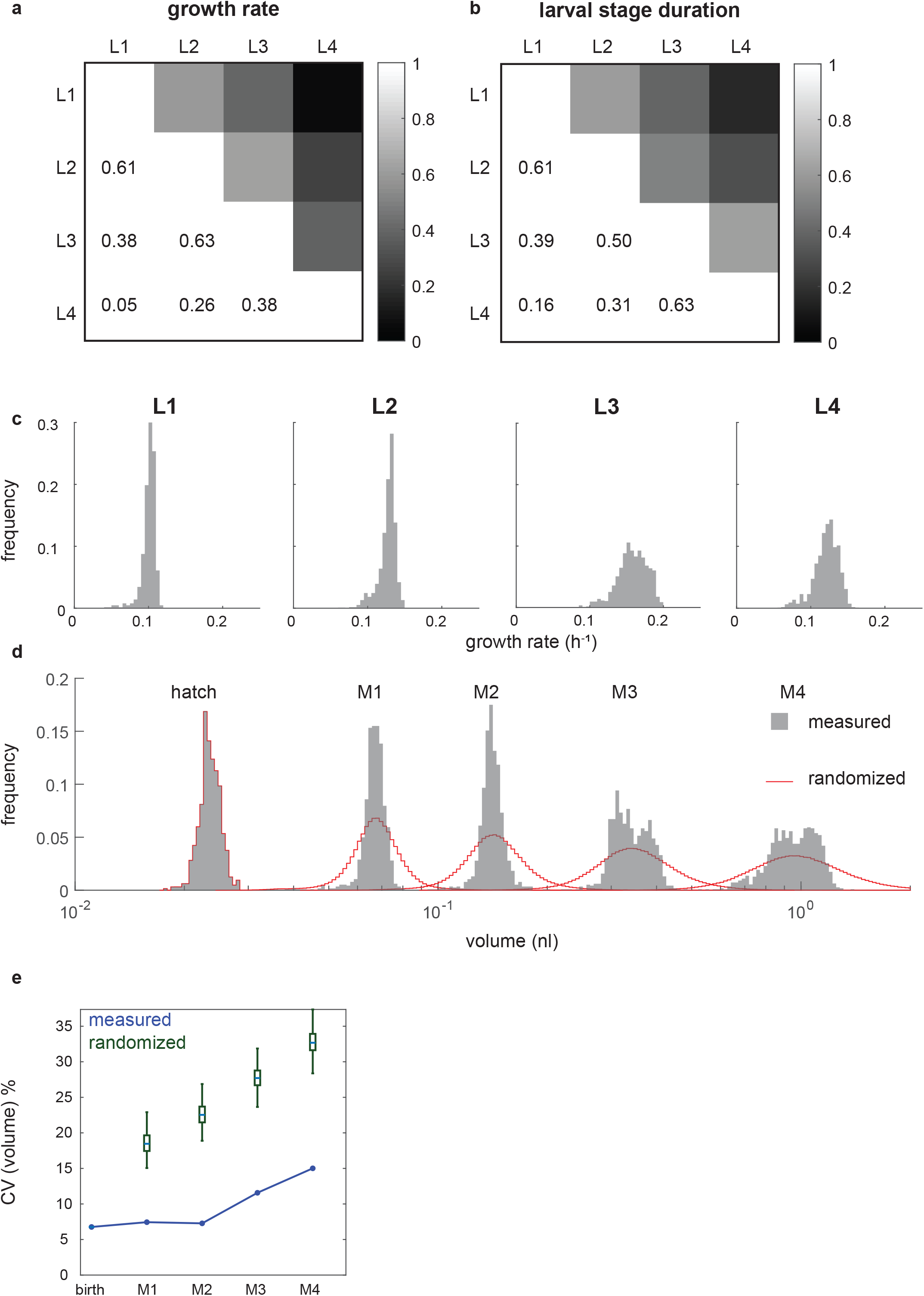
Measurement of body volume divergence without normalization. *(relates to Figure 2)* a, b. Pearson correlation coefficients of growth rates and larval stage duration between larval stages. c. Histogram of growth rate distribution without normalization to mean of day-to-day repeat. d. Histogram of volume distribution without normalization to mean of day-to-day repeat. Red line indicates volume divergence expected by random shuffling of growth rates and duration of larval stages. e. CV of volume for the four larval moults without normalization to mean of day-to-day repeat. Blue line shows measured CV. Green: distribution of randomly shuffled controls for 1000 iterations of random shuffling.

**Figure S3.**
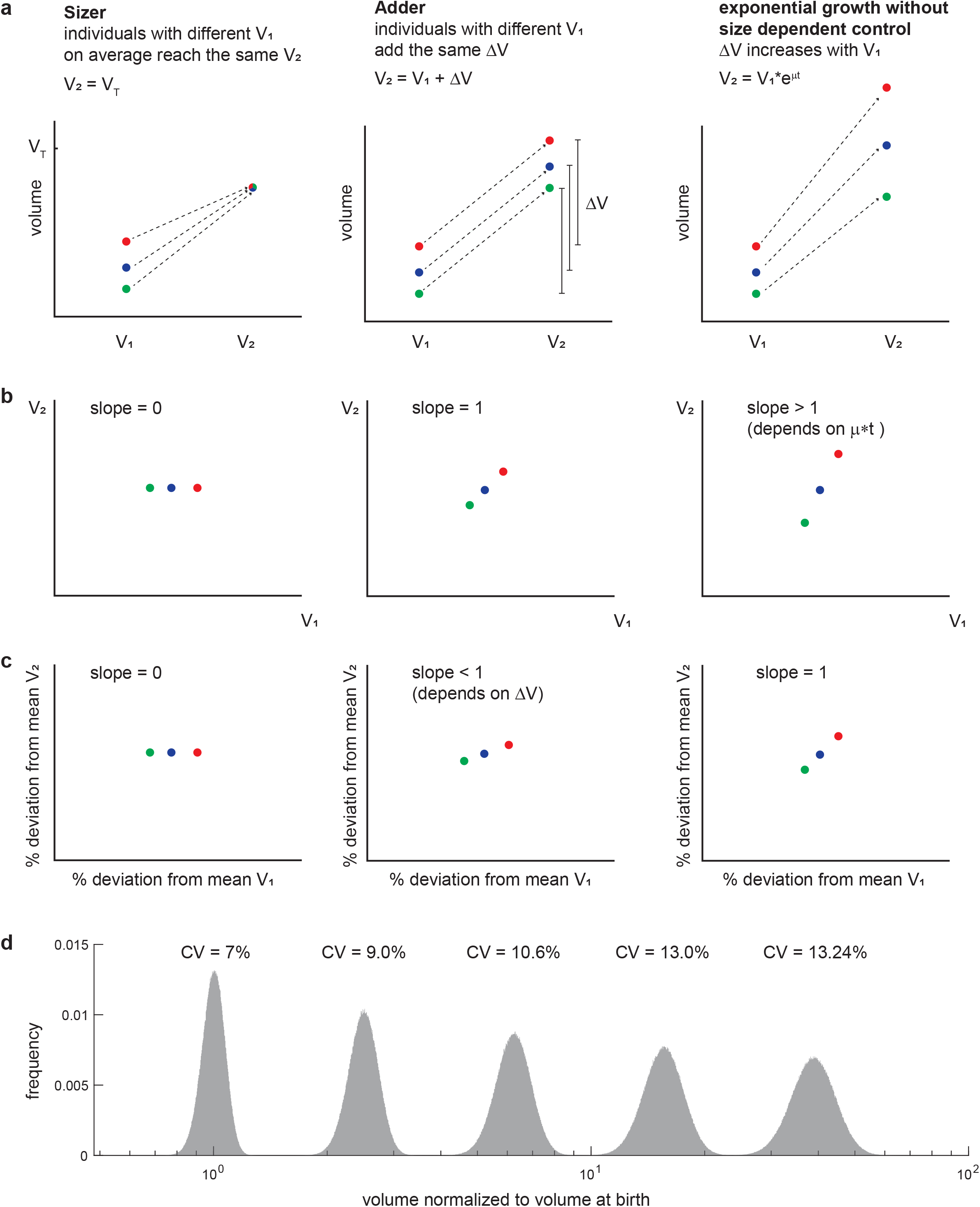
Schematic Illustration of adder, sizer, and folder models. *(related to Figure 3)* a. Schematic representation of 3 individuals that differ in their size at larval stage entry (V_1_). For a sizer, V_2_ is independent of V_1_. For an adder, the size difference ΔV = V_2_ – V_1_ is independent of V_1._ For size-independent exponential growth, ΔV scales positively with V_1_. b. Expected relations between V_1_ and V_2_ for sizers, adders, and size-independent exponential growth. c. Expected relations between V_1_ and V_2_ for sizers, adders, and size-independent exponential growth after normalization to the population mean. d. Simulation of volume divergence of a hypothetical worm population to illustrate imperfectness of a folder in preventing volume divergence. For the simulation, the volume at birth of 100 ‘000 individuals was randomly drawn from a normal distribution with mean = 1 and standard deviation = 0.07. Subsequently, the volume of the following larval stage iteratively was computed by a fold change drawn randomly from a normal distribution with mean = 2.5 and standard deviation = 0.14.

**Figure S4.**
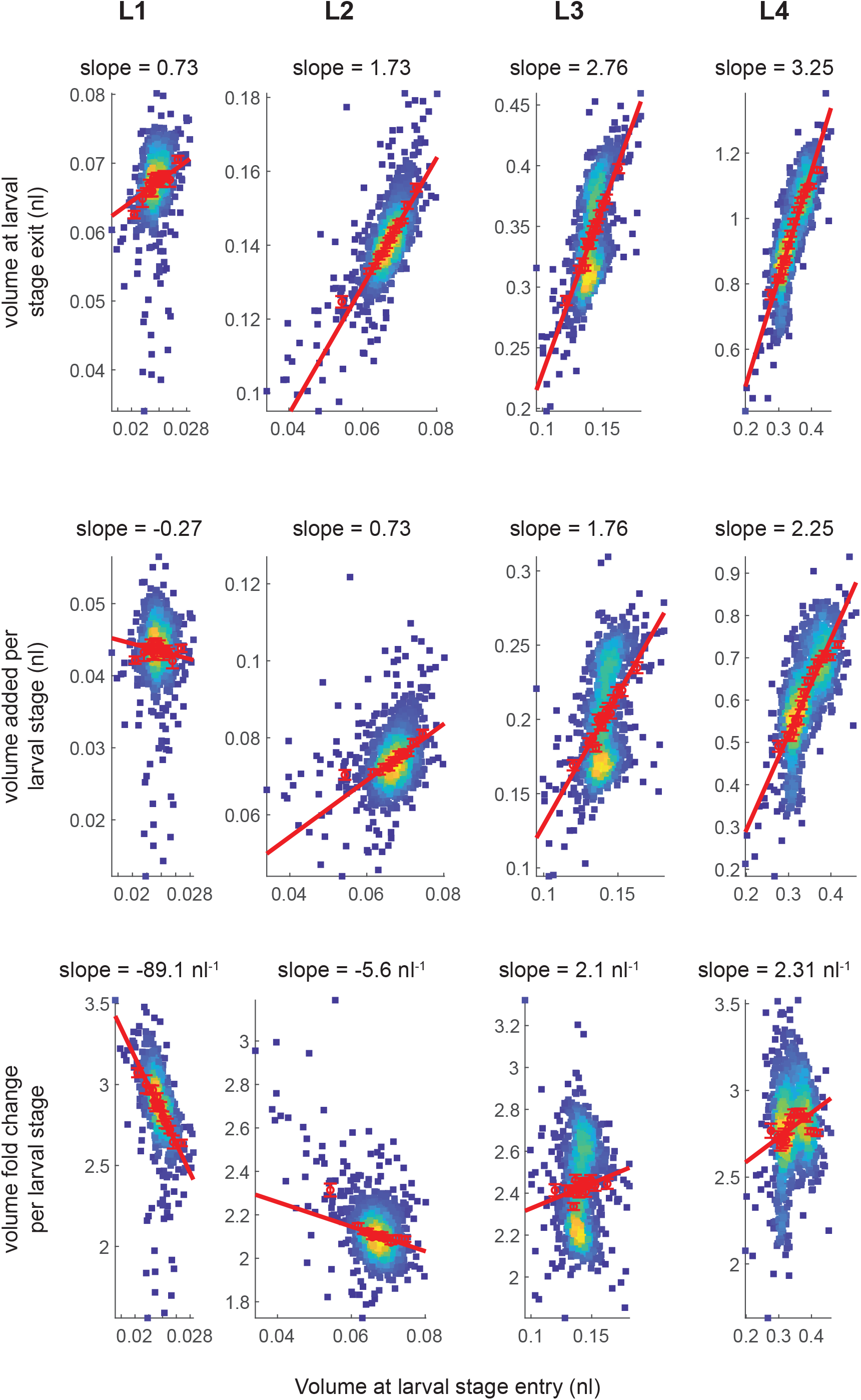
adder, sizer, folder models for unnormalized data. *(related to Figure 3)* Scatter plot of the volumes at larval stage entry and larval stage exit (top), the absolute volumes added per larval stage (middle), and volume fold changes (bottom). Data was not normalized to mean of day-to-day repeat. Red circles: moving average along the x-Axis +/- SEM. Red trendline: regression to moving average excluding the first and the last bin.

**Figure S5.**
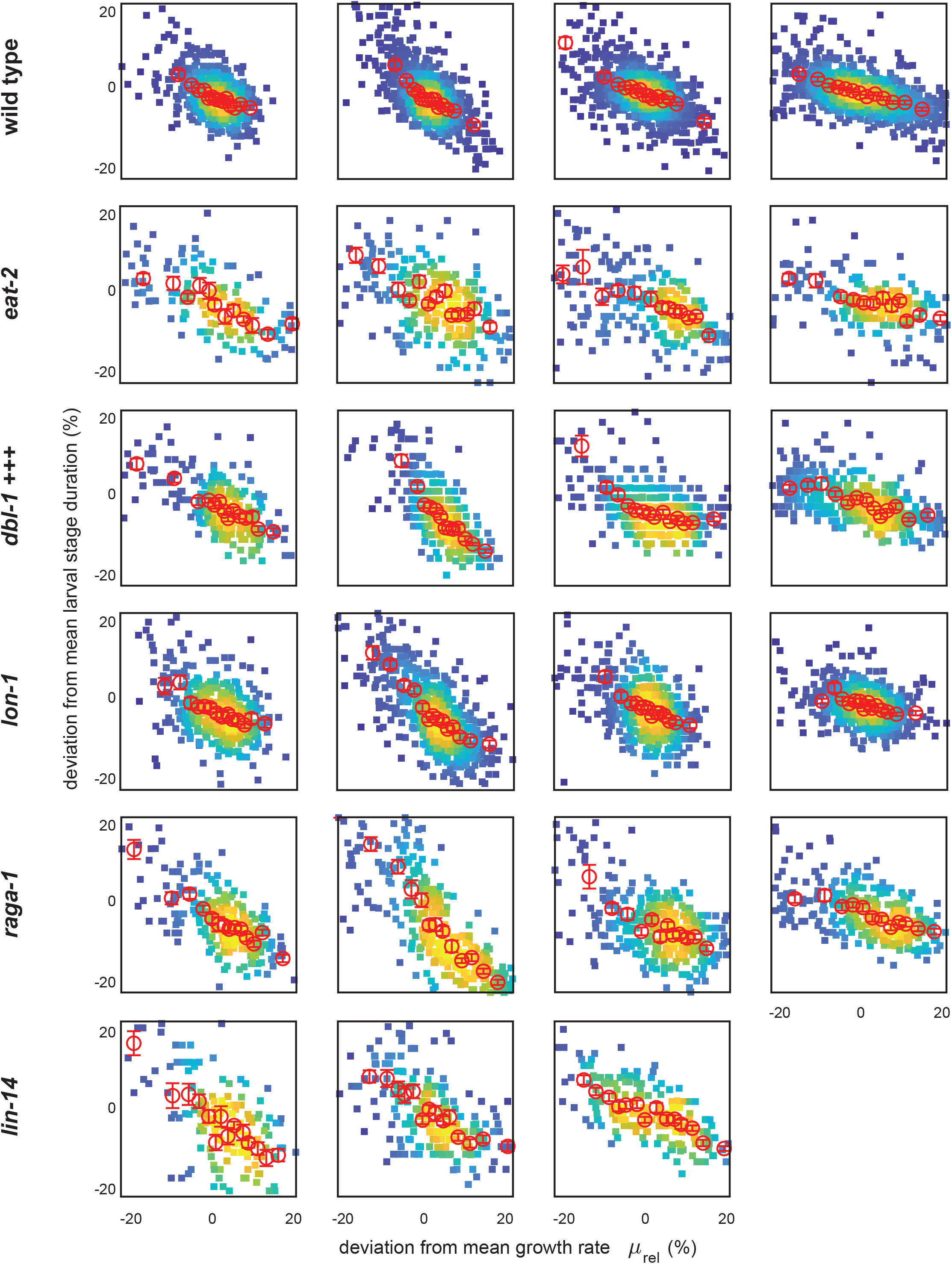
Growth rate and larval stage duration remain anti-correlated in mutant strains. *(related to Figure 4)* Scatter plot of growth rate vs. larval stage duration shown as % deviation from the mean for indicated mutants. Colour indicates point density. Red circles are moving average along x-Axis +/- SEM.

**Figure S6.**
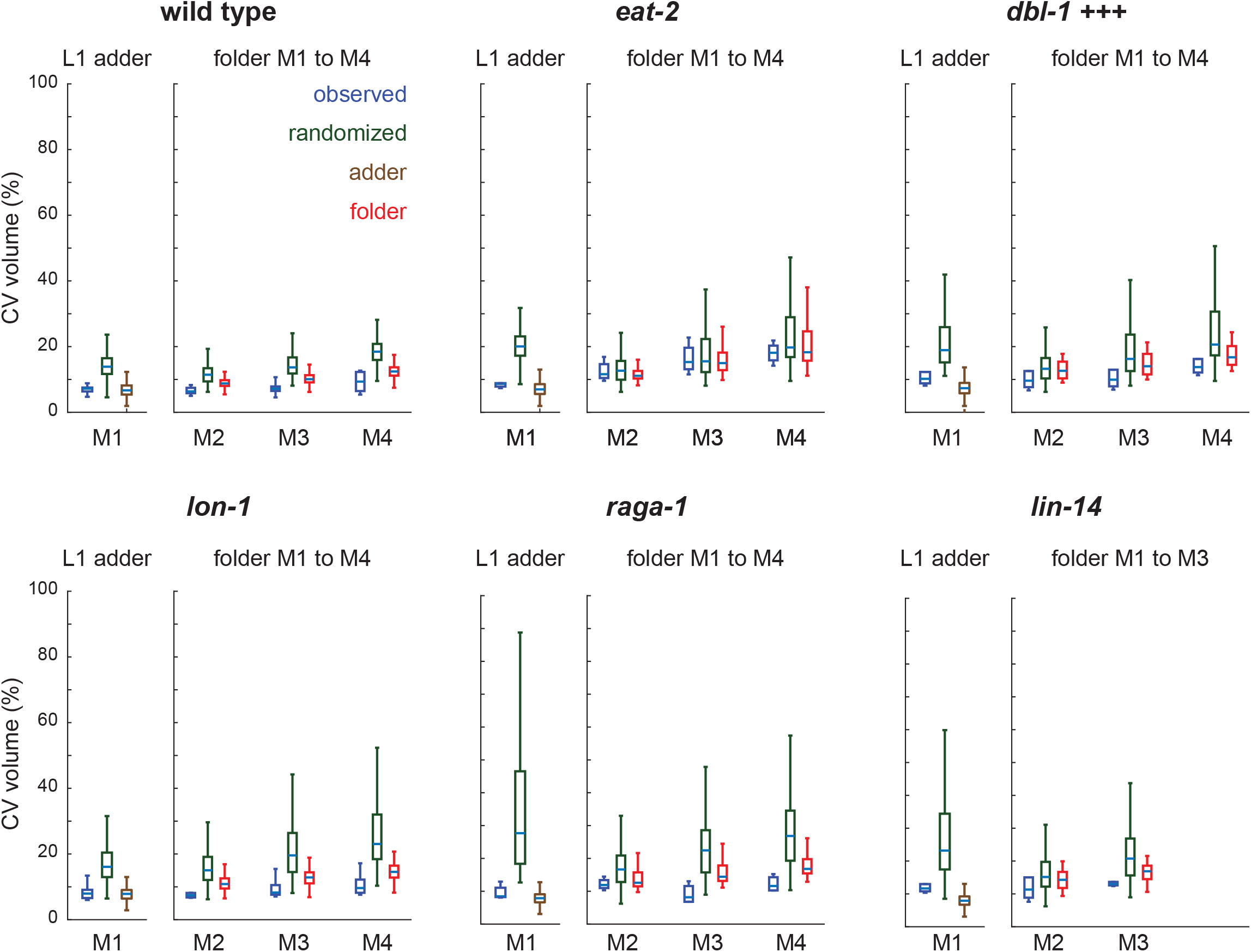
Mutants with perturbed growth rate do not diverge more than expected in volume. *(related to Figure 4)* Comparison of CV of volume to adder and folder models for indicated mutant backgrounds. Blue: box plot of the average CV measured experimentally in the different day-to-day repeats. Green: expected CV based on 1000x random shuffling of growth rate and larval stage duration. Brown: CV expected from adder model during L1. Red: CV expected from folder model during L2 to L4. Randomizations was done separately for L1 and for L2 to L4 starting from measured volume distributions at birth and at M1, respectively.

**Figure S7.**
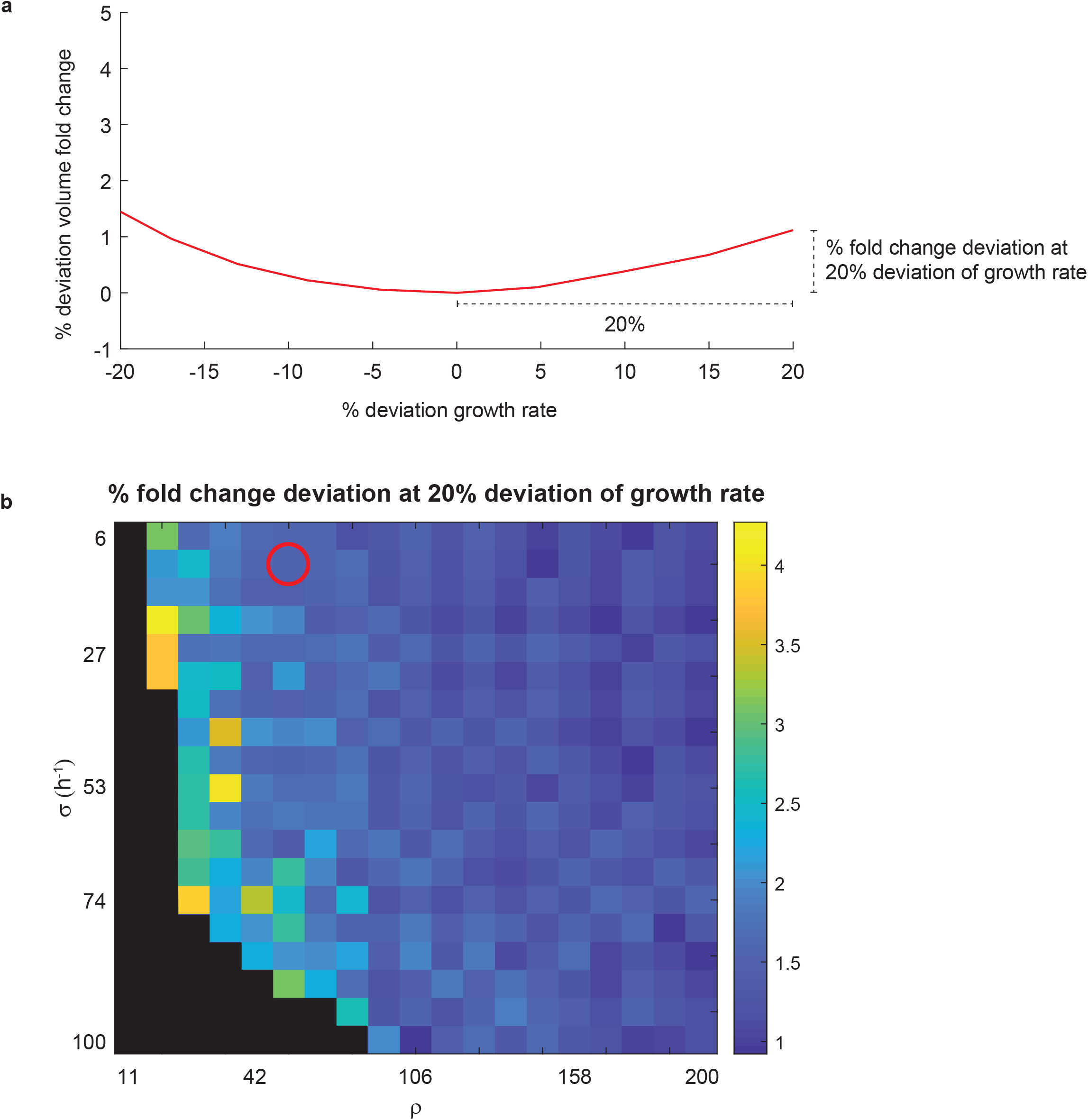
Model behaviour is robust to changes of parameter values. *(related to Figure 5)* a. Illustration of the deviation in fold change plotted in (b). The volume fold change in the model was computed for a range of parameter values for ρ and σ as a function of μ. For each pair of ρ and σ the growth rate μ_0_ was determined, for which the fold change, and hence the first derivative of the fold μ change with respect to μ was minimized (μ_0_ is the μ, at which 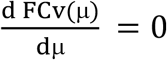). Subsequently, the relative change in fold change after increasing or decreasing μ by 20% was computed. (b) shows the larger one of the two values. b. Volume fold change deviation from reference at a 20% change of μ for a range of model parameters. Deviation is less than 2% from for a large parameter space. Black areas are parameter values for which the system does not oscillate. Red circle indicates parameters used in Fig 5c-f.

## References

1. Blanckenhorn, W. U. The Evolution of Body Size: What Keeps Organisms Small? The Quarterly Review of Biology 75, 385–407 (2000).

2. Gokhale, R. H. & Shingleton, A. W. Size control: the developmental physiology of body and organ size regulation. WIREs Developmental Biology 4, 335–356 (2015).

3. Savage-Dunn, C. & Padgett, R. W. The TGF-β Family in Caenorhabditis elegans. Cold Spring Harb Perspect Biol 9, (2017).

4. Nijhout, H. F. & Williams, C. M. Control of Moulting and Metamorphosis in the Tobacco Hornworm, Manduca Sexta (L.): Growth of the Last-Instar Larva and the Decision to Pupate. Journal of Experimental Biology 61, 481–491 (1974).

5. Uppaluri, S. & Brangwynne, C. P. A size threshold governs Caenorhabditis elegans developmental progression. Proceedings of the Royal Society B: Biological Sciences 282, (2015).

6. Baron, J. et al. Catch-up growth after glucocorticoid excess: a mechanism intrinsic to the growth plate. Endocrinology 135, 1367–1371 (1994).

7. Tanner, J. M. Regulation of Growth in Size in Mammals. Nature 199, 845–850 (1963).

8. Perez, M. F., Francesconi, M., Hidalgo-Carcedo, C. & Lehner, B. Maternal age generates phenotypic variation in Caenorhabditis elegans. Nature 552, 106 (2017).

9. Kinser, H. E., Mosley, M. C., Plutzer, I. B. & Pincus, Z. Global, cell non-autonomous gene regulation drives individual lifespan among isogenic C. elegans. Elife 10, (2021).

10. Burnaevskiy, N. et al. Chaperone biomarkers of lifespan and penetrance track the dosages of many other proteins. Nat Commun 10, 5725 (2019).

11. Gritti, N., Kienle, S., Filina, O. & van Zon, J. S. Long-term time-lapse microscopy of C. elegans post-embryonic development. Nature Communications 7, 12500 (2016).

12. Campos, M. et al. A Constant Size Extension Drives Bacterial Cell Size Homeostasis. Cell 159, 1433–1446 (2014).

13. Soifer, I., Robert, L. & Amir, A. Single-Cell Analysis of Growth in Budding Yeast and Bacteria Reveals a Common Size Regulation Strategy. Current Biology 26, 356–361 (2016).

14. Wallden, M., Fange, D., Lundius, E. G., Baltekin, Ö. & Elf, J. The Synchronization of Replication and Division Cycles in Individual E. coli Cells. Cell 166, 729–739 (2016).

15. Witz, G., van Nimwegen, E. & Julou, T. Initiation of chromosome replication controls both division and replication cycles in E. coli through a double-adder mechanism. eLife 8, e48063 (2019).

16. Taheri-Araghi, S. et al. Cell-Size Control and Homeostasis in Bacteria. Current Biology 25, 385– 391 (2015).

17. Xie, S. & Skotheim, J. M. A G1 Sizer Coordinates Growth and Division in the Mouse Epidermis. Current Biology 30, 916-924.e2 (2020).

18. Fantes, P. A. Control of cell size and cycle time in Schizosaccharomyces pombe. Journal of Cell Science 24, 51–67 (1977).

19. Chandler-Brown, D., Schmoller, K. M., Winetraub, Y. & Skotheim, J. M. The Adder Phenomenon Emerges from Independent Control of Pre- and Post-Start Phases of the Budding Yeast Cell Cycle. Current Biology 27, 2774-2783.e3 (2017).

20. Yu, F. B. et al. Long-term microfluidic tracking of coccoid cyanobacterial cells reveals robust control of division timing. BMC Biol 15, 11 (2017).

21. Deforet, M., van Ditmarsch, D. & Xavier, J. B. Cell-Size Homeostasis and the Incremental Rule in a Bacterial Pathogen. Biophysical Journal 109, 521–528 (2015).

22. Turek, M., Besseling, J. & Bringmann, H. Agarose Microchambers for Long-term Calcium Imaging of Caenorhabditis elegans. JoVE e52742 (2015) doi:10.3791/52742.

23. Meeuse, M. W. et al. Developmental function and state transitions of a gene expression oscillator in Caenorhabditis elegans. Mol Syst Biol 16, (2020).

24. Keil, W., Kutscher, L. M., Shaham, S. & Siggia, E. D. Long-Term High-Resolution Imaging of Developing C. elegans Larvae with Microfluidics. Developmental Cell 40, 202–214 (2017).

25. Zhang, W. B. et al. Extended Twilight among Isogenic C. elegans Causes a Disproportionate Scaling between Lifespan and Health. Cell Systems 3, 333-345.e4 (2016).

26. Sulston, J. E. & Horvitz, H. R. Post-embryonic cell lineages of the nematode, Caenorhabditis elegans. Developmental Biology 56, 110–156 (1977).

27. Hendriks, G. J., Gaidatzis, D., Aeschimann, F. & Grosshans, H. Extensive oscillatory gene expression during C. elegans larval development. Molecular cell 53, 380–92 (2014).

28. Kim, D. H., Grün, D. & van Oudenaarden, A. Dampening of expression oscillations by synchronous regulation of a microRNA and its target. Nature genetics 45, 1337–44 (2013).

29. Moore, B. T., Jordan, J. M. & Baugh, L. R. WormSizer: High-throughput Analysis of Nematode Size and Shape. PLOS ONE 8, e57142 (2013).

30. Singh, R. N. & Sulston, J. E. Some Observations On Moulting in Caenorhabditis Elegans. Nematologica 24, 63–71 (1978).

31. Olmedo, M., Geibel, M., Artal-Sanz, M. & Merrow, M. A High-Throughput Method for the Analysis of Larval Developmental Phenotypes in Caenorhabditis elegans. Genetics 201, 443–448 (2015).

32. Faerberg, D. F., Gurarie, V. & Ruvinsky, I. Inferring temporal organization of postembryonic development from high-content behavioral tracking. Developmental Biology 475, 54–64 (2021).

33. Knight, C. G., Patel, M. N., Azevedo, R. B. R. & Leroi, A. M. A novel mode of ecdysozoan growth in Caenorhabditis elegans. Evol. Dev. 4, 16–27 (2002).

34. Godin, M. et al. Using buoyant mass to measure the growth of single cells. Nature Methods 7, 387–390 (2010).

35. Lakowski, B. & Hekimi, S. The genetics of caloric restriction in Caenorhabditis elegans. Proceedings of the National Academy of Sciences 95, 13091–13096 (1998).

36. Schreiber, M. A., Pierce-Shimomura, J. T., Chan, S., Parry, D. & McIntire, S. L. Manipulation of Behavioral Decline in Caenorhabditis elegans with the Rag GTPase raga-1. PLOS Genetics 6, e1000972 (2010).

37. Sancak, Y. et al. The Rag GTPases Bind Raptor and Mediate Amino Acid Signaling to mTORC1. Science 320, 1496–1501 (2008).

38. Suzuki, Y. et al. A BMP homolog acts as a dose-dependent regulator of body size and male tail patterning in Caenorhabditis elegans. Development 126, 241–250 (1999).

39. Brenner, S. THE GENETICS OF CAENORHABDITIS ELEGANS. Genetics 77, 71–94 (1974).

40. Ambros, V. & Horvitz, H. R. Heterochronic mutants of the nematode Caenorhabditis elegans. Science 226, 409–416 (1984).

41. Maduzia, L. L. et al. lon-1 Regulates Caenorhabditis elegans Body Size Downstream of the dbl-1 TGFβ Signaling Pathway. Developmental Biology 246, 418–428 (2002).

42. Novak, B. & Tyson, J. J. Design Principles of Biochemical Oscillators. Nat Rev Mol Cell Biol 9, 981–991 (2008).

43. Guantes, R. & Poyatos, J. F. Dynamical Principles of Two-Component Genetic Oscillators. PLOS Computational Biology 2, e30 (2006).

44. Alon, U. An introduction to systems biology: design principles of biological circuits. (CRC press, 2006).

45. Conrad, E., Mayo, A. E., Ninfa, A. J. & Forger, D. B. Rate constants rather than biochemical mechanism determine behaviour of genetic clocks. Journal of The Royal Society Interface 5, S9– S15 (2008).

46. Filina, O., Haagmans, R. & Zon, J. S. van. Temporal scaling in C. elegans larval development. bioRxiv 2020.09.21.306423 (2020) doi:10.1101/2020.09.21.306423.

47. Scott, M., Gunderson, C. W., Mateescu, E. M., Zhang, Z. & Hwa, T. Interdependence of cell growth and gene expression: origins and consequences. Science (New York, N.Y.) 330, 1099–102 (2010).

48. Towbin, B. D. et al. Optimality and sub-optimality in a bacterial growth law. Nature Communications 8, 14123 (2017).

49. Goodman, M. B. & Sengupta, P. How Caenorhabditis elegans Senses Mechanical Stress, Temperature, and Other Physical Stimuli. Genetics 212, 25–51 (2019).

50. Rensing, L., Meyer-Grahle, U. & Ruoff, P. Biological Timing and the Clock Metaphor: Oscillatory and Hourglass Mechanisms. Chronobiology International 18, 329–369 (2001).

51. Oates, A. C., Morelli, L. G. & Ares, S. Patterning embryos with oscillations: structure, function and dynamics of the vertebrate segmentation clock. Development 139, 625–639 (2012).

52. Frøkjaer-Jensen, C. et al. Single-copy insertion of transgenes in Caenorhabditis elegans. Nat Genet 40, 1375–1383 (2008).

